# Quantification of durable CRISPR-based gene silencing activity

**DOI:** 10.1101/2021.03.31.436355

**Authors:** Muneaki Nakamura, Alexis Ivec, Yuchen Gao, Lei S. Qi

**Affiliations:** Department of Bioengineering, Stanford, CA 94305, United States; Program in Human Biology, Stanford, CA 94305, United States; Cancer Biology Program, Stanford, CA 94305, United States; Department of Chemical and Systems Biology, Stanford, CA 94305, United States; ChEM-H Institute, Stanford, CA 94305, United States; Mammoth Biosciences, South San Francisco, CA 94080, United States

## Abstract

Development of CRISPR-based technologies for regulating gene expression stands to provide novel methods for the study and engineering of biological behavior. New tools capable of inducing long-lasting changes in gene expression will increase the utility of these techniques, providing durable effects from one-time doses of reagents. We describe here a reporter system for quantifying the ability of CRISPR-based effectors to induce stable gene repression. We observe a continuous gradation of the ability of these effectors to silence gene expression, depending on the domain composition and configuration. We also report the creation of a single CRISPR protein capable of producing durable gene silencing. This assay should allow for the continued development of enhanced gene repression tools which will be useful in a wide array of biological research and engineering applications.

## Introduction

The adoption of CRISPR systems has enabled a wide range of synthetic biology applications by allowing the rapid targeting of CRISPR-associated (Cas) proteins to almost anywhere in the genome via a guide RNA (gRNA). In particular, nuclease-deactivated Cas (dCas) proteins have been applied for use of control of gene expression via directed dCas binding^1^ and the localization of domains driving gene up- and down-regulation (termed CRISPRa and CRISPRi, respectively)^2,3^. A typical feature of CRISPRi/a technologies is that gene expression changes are maintained only while the effector domains remain actively targeted to the locus of interest^4^. Although this may be beneficial in many contexts, in other applications, it may be desirable to generate gene regulation changes that can persist after a transient dose of CRISPR effector.

The co-option of epigenetic processes has long been examined as a possible route for inducing long-lasting gene expression changes^5^, with much research in particular focusing on DNA CpG methylation as a mark highly associated with gene silencing. However, the gene expression changes induced by engineered DNA methylation remain quite modest. A report describing the simultaneous use of the Krüppel associated box (KRAB) domain commonly used in CRISPRi along with DNMT3A and DNMT3L domains (involved in *de novo* DNA methylation) showed significant, long-lasting gene repression^6^. This study, along with others using the same combination of domains^7,8^, however, demonstrated mixed repression capability depending on the gene target, raising the question of the context-dependence of this approach. To address this issue, we endeavored to create a synthetic reporter system in order to systematically assess the ability of domains to produce stable gene silencing. As a test case, we used this reporter system to demonstrate the creation of a single dCas protein effector capable of inducing durable gene silencing.

## Materials and Methods

### Experimental Design

Our experimental design was broken down into the following portions. First, we created a new standardized reporter cell line that could be used to quantitatively compare the activity of CRISPR-based effectors. Once created, we transiently introduced CRISPR effectors into this reporter line and monitored the resulting change on reporter gene expression over time. Analysis of these results informed the design of next generation effector combinations.

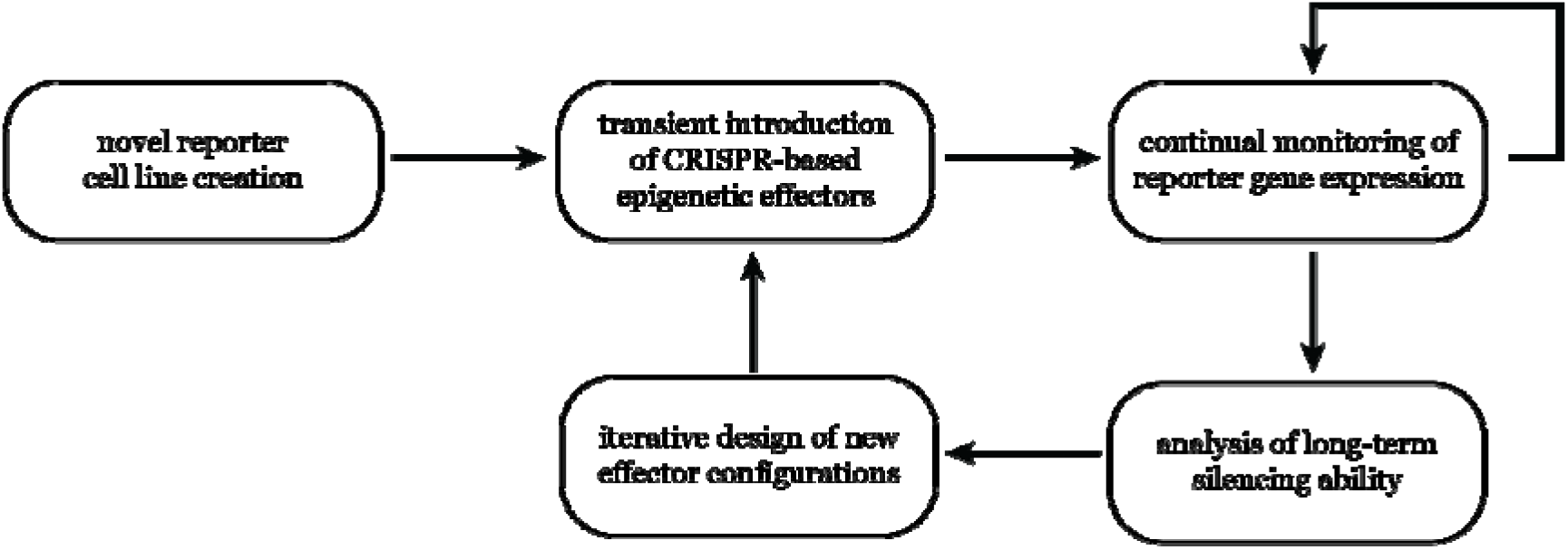

### Cell culture

HEK293T cells were cultured in DMEM containing sodium pyruvate, GlutaMAX, and 4.5 g / L glucose (Thermo Fisher Scientific) supplemented with 10 % FBS. Cells were not tested for mycoplasma contamination.

HEK293T cells bearing EGFP expressed from the SV40 promoter, used in previous studies^4,9^, were lentivirally transduced with a cassette expressing sgRNA targeting the SV40 promoter and BFP and puromycin resistance proteins. Selection with 2 μg / mL puromycin resulted in near purity of cells bearing both cassettes.

Transfection of constructs was performed in 12-well plates using TransIT-LT1 reagent (Mirus) with 1 μg DNA split evenly among plasmids used in each transfection. Cells were exposed to 400 μg / mL Zeocin (Thermo Fisher Scientific) days 2-5 post-transfection. Cells were serially passaged and assessed for fluorescence via flow cytometry with a CytoFLEX S machine (Beckman Coulter).

### Data analysis

Raw flow cytometry data were compensated and analyzed using FlowJo software, gating for live singlet cells. The resulting data were processed and analyzed using custom scripts.

The data were normalized at each timepoint using a wild-type HEK293T control lacking fluorescent reporter to parameterize a cutoff of negative fluorescence defined at the 99th percentile of GFP signal, scaling GFP fluorescence over all conditions to this value. Timecourses of median fluorescence of each condition were fit using least-squares to an asymmetric Gaussian of form

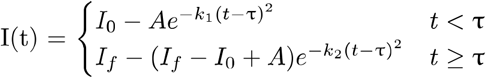

with I indicating fluorescent intensity. I_0_ was parameterized by the fluorescence level of untransfected reporter, and all other parameters were varied to fit the model. Fold-change in intensity was calculated by the ratio I_0_ / I_f_.

Histograms of cell fluorescence distributions were fit to a sum of three Gaussians. The amplitude and mean were allowed to vary, along with a Gaussian width parameter (shared among the three Gaussians).

## Results

### Characterization of the fluorescent reporter system

We based our reporter on a cassette containing GFP driven by the constitutive SV40 promoter, along with a corresponding cassette expressing the relevant gRNA targeting the GFP promoter. These constructs were lentivirally introduced into HEK293T cells. The resultant cells contain all components except the relevant CRISPR protein effectors, allowing the transient introduction of effectors to assess the ability to drive GFP silencing under a consistent cellular environment (Fig. 1a, Supp. Fig. 1a,b).

**Figure 1.**
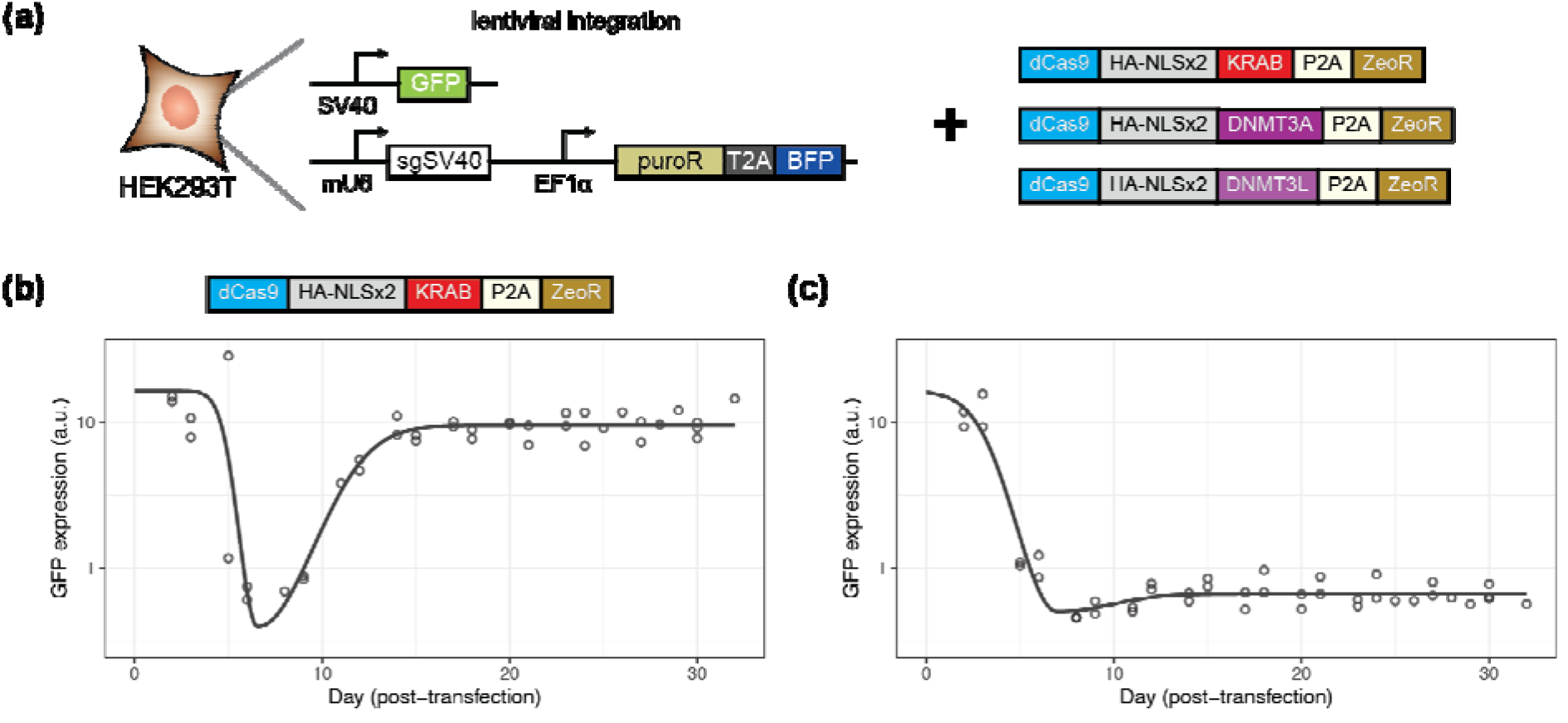
Assay for assessing long-term gene silencing. **(a)** A fluorescent GFP reporter driven by the SV40 promoter is integrated into HEK293T cells, along with a cassette that guides dCas9 to bind to the SV40 promoter. These cells can be transiently transfected with dCas9 fused to various domains to assess the ability to repress GFP over time. **(b-c)** GFP expression over time following transfection of effectors. Dots indicate individual timepoints and model fit is indicated by solid lines. **(b)** Transfection of dCas9-KRAB. **(c)** Co-transfection of dCas9-KRAB, dCas9-DNMT3A, and dCas9-DNMT3L.

We created a set of constructs each bearing one of the three relevant domains (KRAB, DNMT3A, and DNMT3L) fused to *Spy* dCas9 protein and additionally expressing a separate Zeocin-resistance domain via the self-cleaving P2A peptide tag. Reporter cells were transiently transfected with these constructs, subjected to Zeocin selection to enrich for successful introduction of the transfected construct, and the fluorescence monitored over time via flow cytometry.

Consistent with prior studies^2,4,9^, we observed significant GFP repression with dCas9-KRAB shortly following transfection, with subsequent recovery of expression at long timescales (Fig. 1b, Supp. Fig. 1c). Conversely, with co-transfection of all three dCas9 effectors, we saw the formation of a population of cells demonstrating repression of GFP expression that was stable for weeks post-transfection (Fig. 1c, Supp. Fig. 1d). These cells demonstrated a varied response, with some cells silencing more strongly than others, possibly indicating variable delivery of the three components, since each construct bears the same resistance marker (therefore disallowing the distinguishing of cells with all three constructs versus one or two).

### Analysis of novel domain configurations

We examined this possibility further through the reduction from a three-plasmid system to a two-plasmid system, creating combinatorial dual-domain variants fused at the C-terminus of dCas9 (Fig. 2a), co-transfecting each of these dual-domain constructs with the cognate dCas9 bearing the third effector.

**Figure 2.**
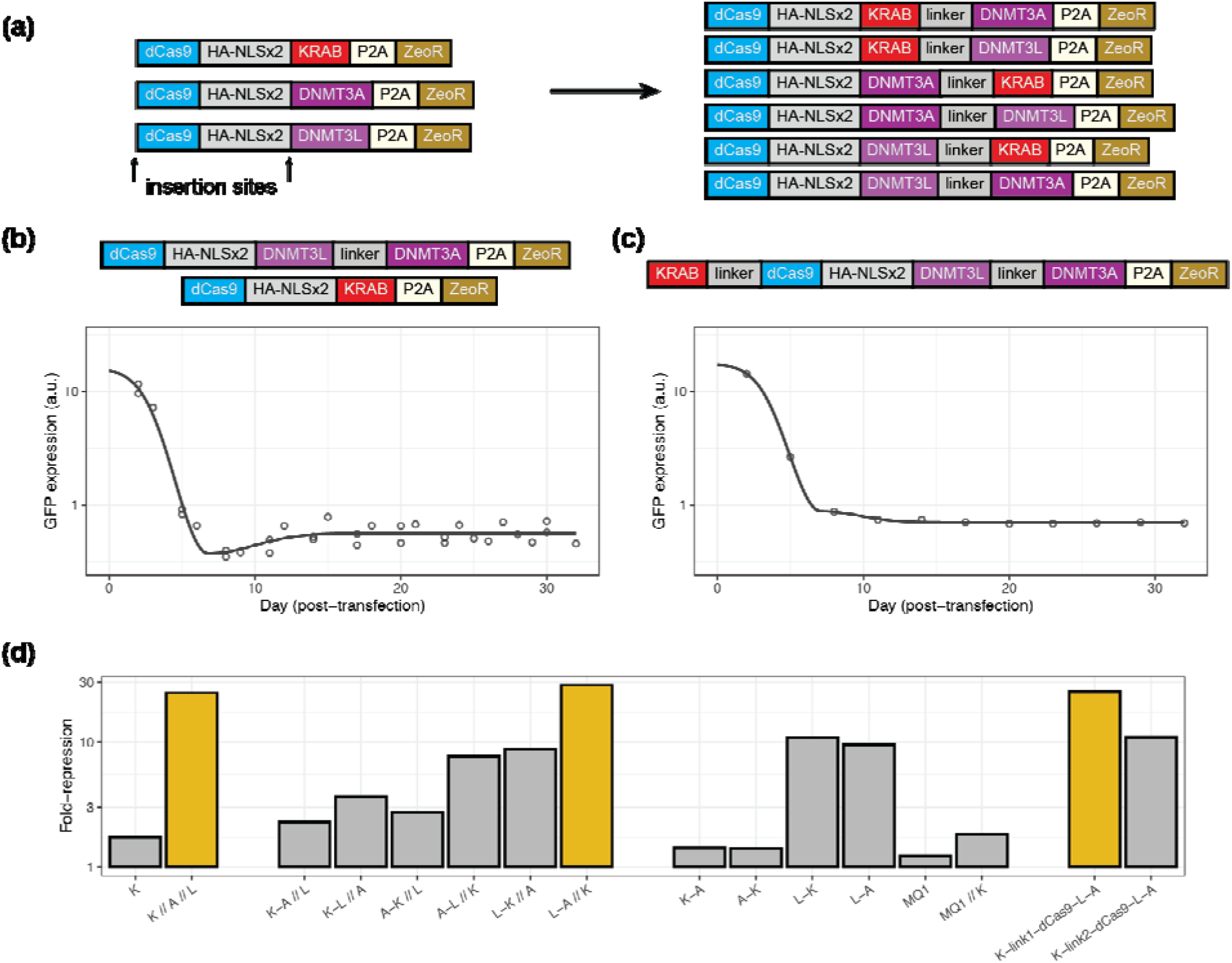
Combinatorial configurations of silencing system. **(a)** Scheme depicting the insertion of combinations of effector domains. The right side indicates combinatorial dual-effector constructs. **(b-c)** GFP expression over time following transfection of effectors. Dots indicate individual timepoints and model fit is indicated by solid lines. **(b)** Co-transfection of dCas9-DNMT3L-DNMT3A + dCas9-KRAB. **(c)** Transfection of KRAB-dCas9-DNMT3L-DNMT3A all-in-one construct. **(d)** Summary of long-term silencing ability of various configurations relative to untransfected cells. All domains are fused to dCas9; where otherwise not indicated, the domain is fused to the C-terminus. K = KRAB, A = DNMT3A, and L = DNMT3L. Domains connected by dashes indicate multiple fusions. Double slashes separate multiple dCas9 fusions that were co-transfected.

We found that the silencing capabilities of these constructs varied widely (Supp. Fig. 2), with many combinations demonstrating impaired long-term repression compared to the three-plasmid system. One combination in particular, using DNMT3L and DNMT3A linked together and co-transfected with KRAB, demonstrated high levels of silencing comparable to the three-plasmid system (Fig. 2b, Supp. Fig. 2f).

Testing the dual-effector constructs alone also revealed differences in silencing ability (Supp. Fig. 3), with constructs containing DNMT3L being more effective at silencing GFP. These results also indicate that stable repression can be achieved without the simultaneous use of all three effector domains. We also tested the ability of a prokaryotic DNA methyltransferase, the MQ1 domain from *M. SssI*^10^ to induce silencing with and without dCas9-KRAB, but observed no silencing effect associated with this domain (Supp. Fig. 4a,b).

We next investigated the possibility of further condensing the domains into a single dCas9 construct. Since all combinations of KRAB and DNMT3A at the C-terminus demonstrated the lowest repressive ability, we hypothesized that unfavorable steric hindrances between the KAP1 complex recruited by KRAB and the DNMT3A:DNMT3L complex may have prevented full activity of the effector domains. Therefore, we used our most active dCas9-DNMT3L-DNMT3A construct and tested N-terminal fusions of the KRAB domain, varying the linker length bridging the KRAB and dCas9. In both cases, we found significant gene silencing (Supp. Fig. 4c,d), with the shorter linker demonstrating large levels of stable repression (Fig. 2c).

Using the data from this reporter assay, we are able to quantitatively assess the level of silencing induced by our various construct combinations. We observe a gradation of silencing capabilities that depends strongly on the domains used and their configuration relative to the dCas9 protein (Fig. 2d). The percentage of cells with negligible GFP expression demonstrated a similar pattern across conditions as the fold-change in expression level (Supp. Fig. 5a).

To further understand the contributors to the silencing process, we examined the cell-to-cell distributions in GFP expression across conditions. We observed that the distributions could be divided into populations of states corresponding to GFP-expressing cells and GFP-negative cells, and that the primary driver of the difference between conditions was the ability of a particular effector combination to drive transitions into the GFP-negative state (Supp. Fig. 5b-d). However, we also observed some contribution from a change of GFP expression within each state, with stronger silencers also lowering the expression across states.

## Discussion

Development of new tools for engineering gene regulation is greatly assisted by assays to quantitatively measure their effect. Our system here enables a simple assay for the distinguishing of domain fusions to dCas9 on the single-cell level without the need for sorting. Using this system, we were able to find a configuration that combines all effectors necessary for inducing long-term silencing onto a single protein. We hypothesize that such a dCas9 effector may be useful in applying epigenetic-based repression to other biological contexts.

Our all-in-one system did not demonstrate markedly greater silencing capability than the three-plasmid system. There are some inherent tradeoffs in reducing the number of constructs – while a higher proportion of the cells may contain all the factors for an all-in-one system, larger construct size results in lower transfection efficiency and lower expression. Whether these factors account for the limit of silencing (in our best constructs, about 60 % of cells achieving total silencing) or whether this reflects limitations in our construct design or assay remains to be determined.

One aspect revealed by our study is the dependence of the silencing activity on the relative configuration of effector domains. KRAB seemed to perform better when given sufficient room in three-dimensional space. Additionally, the better performance of the DNMT3L-DNMT3A fusion relative to the DNMT3A-DNMT3L construct might result from the distances of the N- and C-termini of the DNMT3A and DNMT3L domains when assembled into the active heterotetramer complex^11–13^.

This assay was also able to differentiate between the activity of various methyltransferases. Although the MQ1 domain has been used to apply DNA methylation in mammalian cells^10,14,15^, evidence for its effect on gene expression remains limited^10^. We demonstrate here that it is much less active than DNMT3A / DNMT3L, possibly due to the lack of endogenous epigenetic interactions or its smaller methylation window^10^.

We also demonstrated that significant gene expression changes can be induced using a subset of the domains. Our results are consistent with a prior report that implied a two-state reporter system that can be turned off using KRAB or DNMT3A alone^16^. However, in our assay, we observe a continuum of gene repression levels, with combinations of domains enhancing the transition to a silenced state, hinting a more complex set of states that may play a role in regulating gene expression.

The assay described in this work should allow for the rapid determination of silencing capabilities. By making quantitative measurements of gene expression changes, it should be possible to stringently test novel repression domains and configurations thereof, which will provide enhanced tools for generating long-lasting gene expression changes within cells.

## Author contributions

M.N., L.S.Q., and Y.G. conceived of research. M.N., A.I., and Y.G. provided materials. M.N. and A.I. performed experiments and analyzed data. M.N. wrote the manuscript with input from other authors.

## Supplementary Materials

**Supplementary Figure 1.**
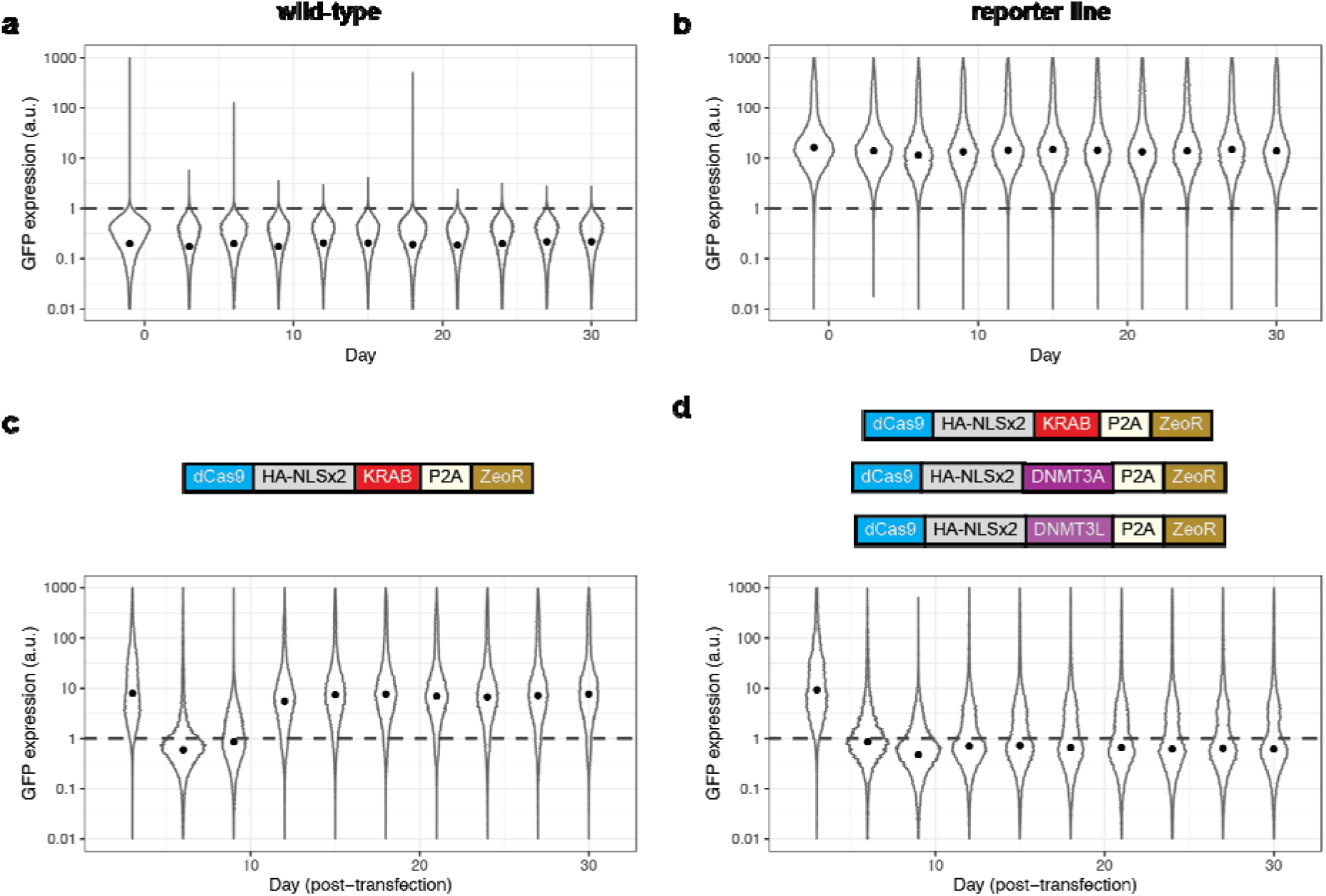
**(a-d)** Representative raw GFP fluorescence distributions over time plotted as violin plots. Median fluorescence indicated by black dot. Dotted line indicates cutoff of no fluorescence, as determined by wild-type HEK293T cells **(a-b)** Traces of cell lines untransfected and not exposed to Zeocin. **(a)** Wild-type HEK293T cell line without integrated reporter. **(b)** Reporter HEK293T cell line with GFP and sgRNA cassette integrated. **(c-d)** Traces of reporter cell line following transfection of indicated dCas9 effector plasmid and Zeocin selection. **(c)** Transfection of dCas9-KRAB alone. **(d)** Co-transfection of dCas9-KRAB, dCas9-DNMT3A, and dCas9-DNMT3L plasmids.

**Supplementary Figure 2.**
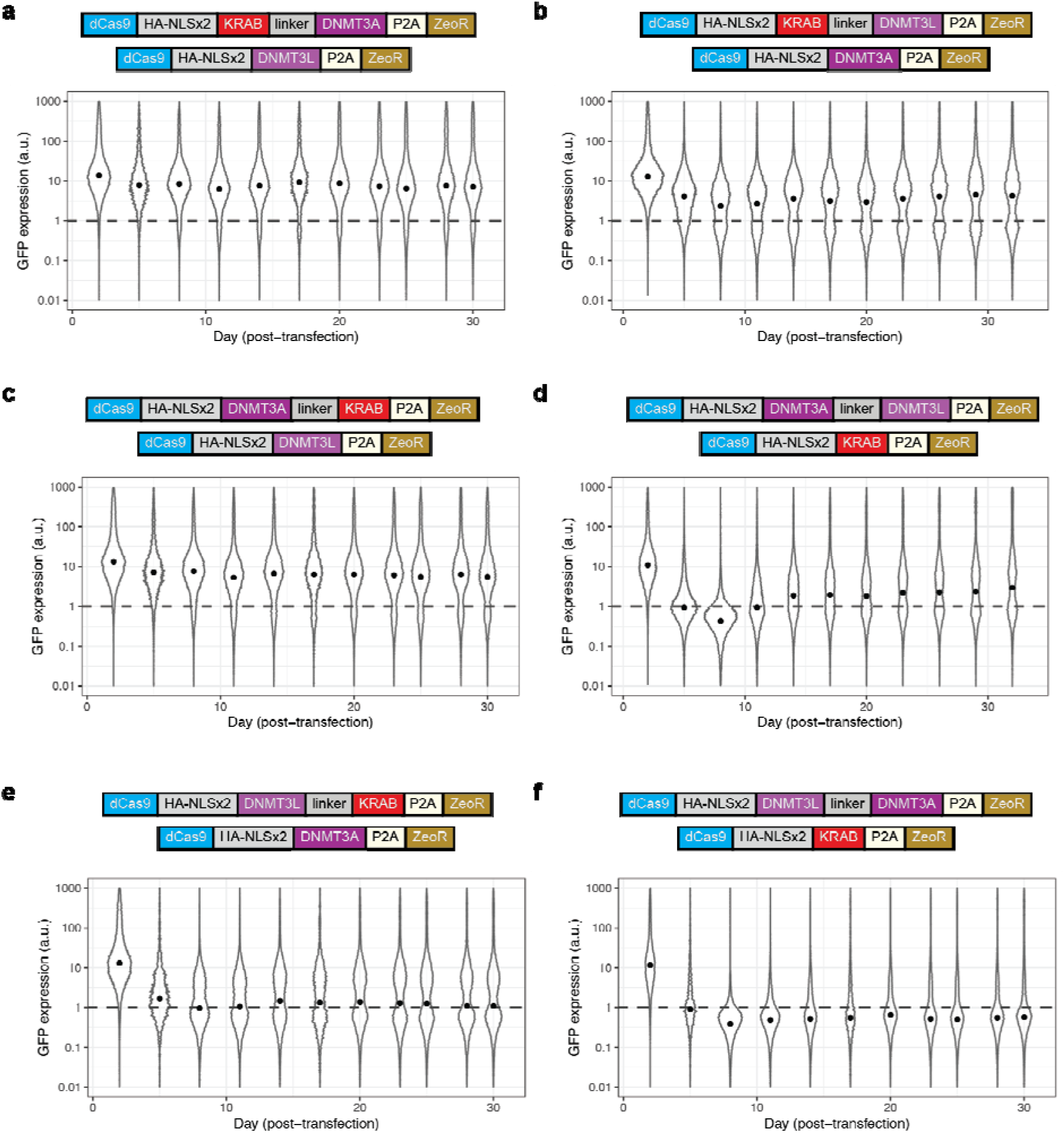
Representative violin plots of raw GFP fluorescence distributions over time following transient transfection and Zeocin selection. Median fluorescence indicated by black dot. Dotted line indicates cut-off of no fluorescence, as determined by wild-type HEK293T cells. **(a)** Co-transfection of dCas9-KRAB-DNMT3A and dCas9-DNMT3L. **(b)** Co-transfection of dCas9-KRAB-DNMT3L and dCas9-DNMT3A. **(c)** Co-transfection of dCas9-DNMT3A-KRAB and dCas9-DNMT3L. **(d)** Co-transfection of dCas9-DNMT3A-DNMT3L and dCas9-KRAB. **(e)** Co-transfection of dCas9-DNMT3L-KRAB and dCas9-DNMT3A. **(f)** Co-transfection of dCas9-DNMT3L-DNMT3A and dCas9-KRAB.

**Supplementary Figure 3.**
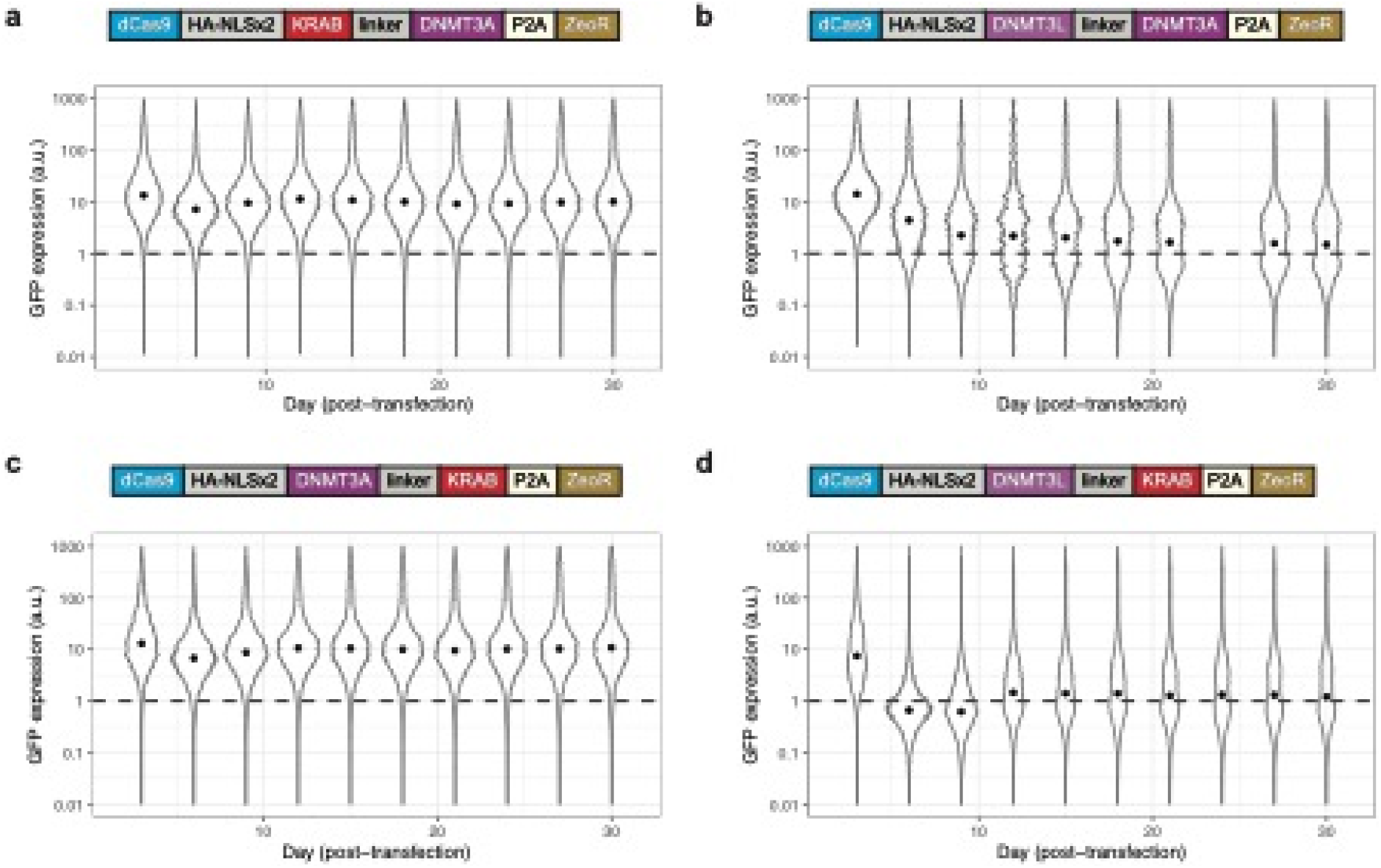
Representative violin plots of raw GFP fluorescence distributions over time following transient transfection and Zeocin selection. Median fluorescence indicated by black dot. Dotted line indicates cut-off of no fluorescence, as determined by wild-type HEK293T cells. **(a)** dCas9-KRAB-DNMT3A. **(b)** dCas9-DNMT3L-DNMT3A. **(c)** dCas9-DNMT3A-KRAB. **(d)** dCas9-DNMT3L-KRAB.

**Supplementary Figure 4.**
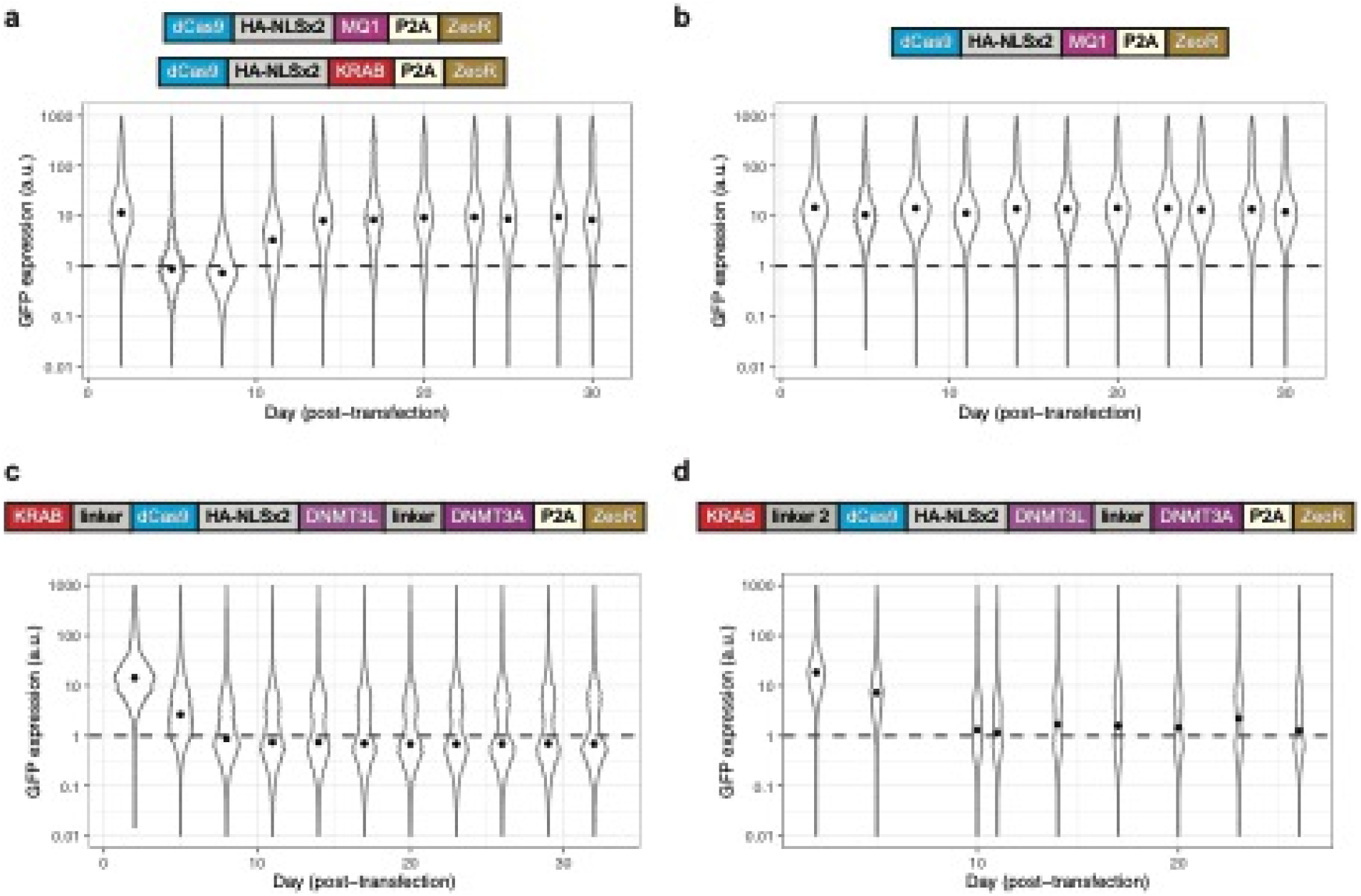
Representative violin plots of raw GFP fluorescence distributions over time following transient transfection and Zeocin selection. Median fluorescence indicated by black dot. Dotted line indicates cut-off of no fluorescence, as determined by wild-type HEK293T cells. **(a)** Co-transfection of dCas9-MQ1 and dCas9-KRAB. **(b)** dCas9-MQ1. **(c)** KRAB-dCas9-DNMT3L-DNMT3A with shorter linker connecting KRAB and dCas9. **(d)** KRAB-dCas9-DNMT3L-DNMT3A with longer linker connecting KRAB and dCas9.

**Supplementary Figure 5.**
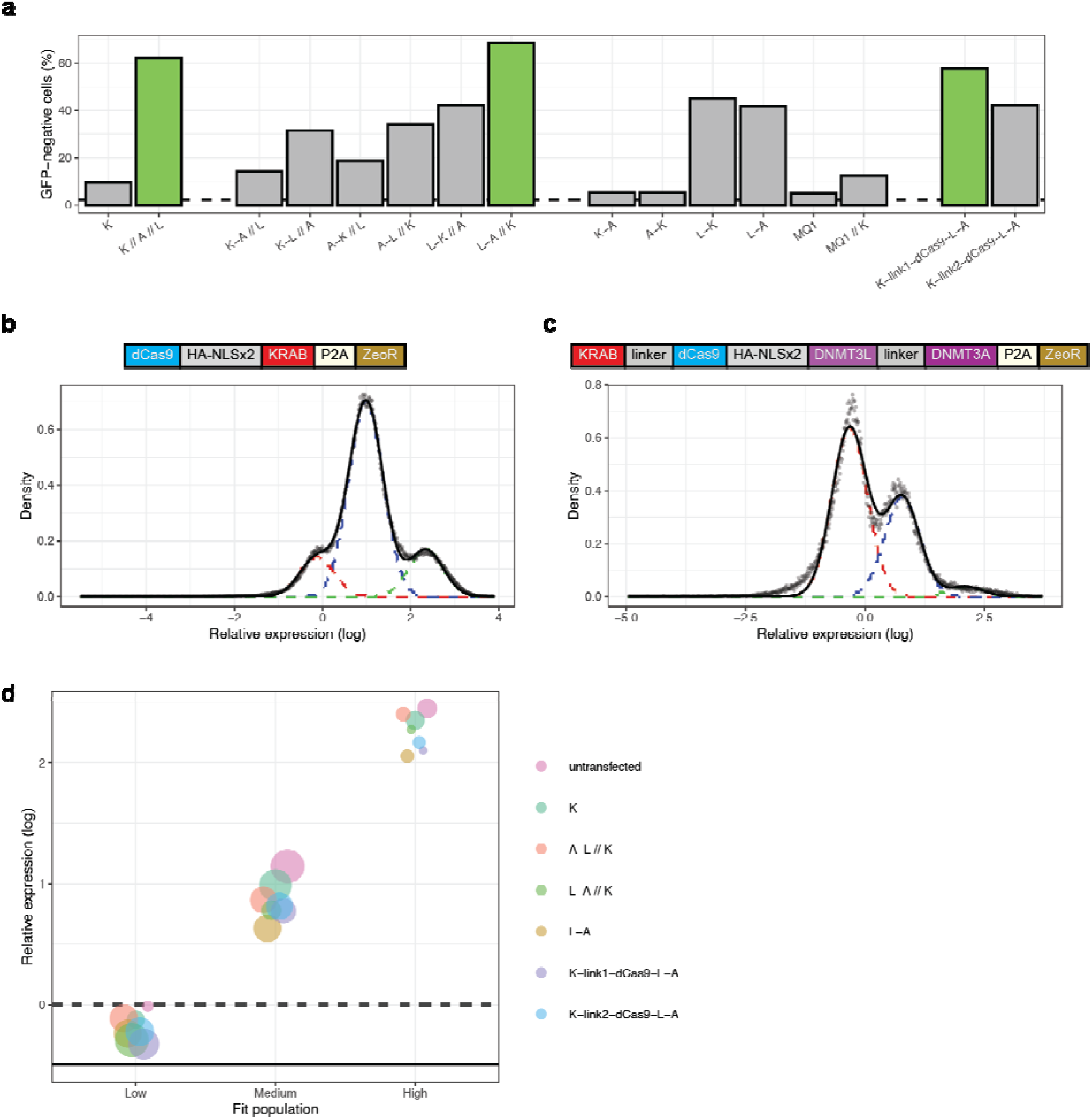
**(a)** Assessment of GFP-negative cells, as determined by cutoff from wild-type cells, across transfection conditions expressed as an average of all measurements more than 21 days post-transfection. All domains are fused to dCas9; where otherwise not indicated, the domain is fused to the C-terminus. K = KRAB, A = DNMT3A, and L = DNMT3L. Domains connected by dashes indicate multiple fusions. Double slashes separate multiple dCas9 fusions that were co-transfected. Dotted line indicates the GFP-negative measurement of un-transfected cells. **(b-c)** Representative fits to histograms of GFP expression (plotted as gray dots). Dashed lines indicate individual peaks of the triple-Gaussian fit, and the black solid line their joint distribution. **(b)** dCas9-KRAB. **(c)** KRAB-dCas9-DNMT3L-DNMT3A with shorter linker connecting KRAB and dCas9. **(d)** Plot of parameters from triple-Gaussian fits for various conditions. Abbreviations as in (a). The low, medium, and high values correspond to the center of the individual Gaussian peaks. The size of the dot corresponds to the relative area of that Gaussian for each condition. The dashed line indicates the negative-fluorescence cutoff. The solid line indicates average of the wild-type (no fluorescence) population.

